# An opaque cell-specific expression program of secreted proteases and transporters allows cell-type cooperation in *Candida albicans*

**DOI:** 10.1101/2020.04.21.053991

**Authors:** Matthew B. Lohse, Lucas R. Brenes, Naomi Ziv, Michael B. Winter, Charles S. Craik, Alexander D. Johnson

**Affiliations:** Department of Microbiology and Immunology, University of California – San Francisco, San Francisco, CA 94143; Department of Pharmaceutical Chemistry, University of California – San Francisco, San Francisco, CA 94143

**Author notes:** Biology Graduate Program, Massachusetts Institute of Technology, Cambridge, MA 02139. CytomX Therapeutics Inc., South San Francisco, CA 94080. To whom correspondence should be addressed: Alexander D. Johnson, Department of Microbiology and Immunology, University of California, San Francisco, 600 16^th^ Street, MBGH Room N372E, Box 2200, San Francisco, CA 94143-2200.

**Keywords:** white-opaque switching, fungal pathogenesis, microbiome, protease

## Abstract

An unusual feature of the opportunistic pathogen *C. albicans* is its ability to stochastically switch between two distinct, heritable cell types called white and opaque. Here, we show that only opaque cells, in response to environmental signals, massively up-regulate a specific group of secreted proteases and peptide transporters, allowing exceptionally efficient use of proteins as sources of nitrogen. We identify the specific proteases (members of the secreted aspartyl protease (*SAP*) family) needed for opaque cells to proliferate under these conditions, and we identify four transcriptional regulators of this specialized proteolysis and uptake program. We also show that, in mixed cultures, opaque cells enable white cells to also proliferate efficiently when proteins are the sole nitrogen source. Based on these observations, we suggest that one role of white-opaque switching is to create mixed populations where the different phenotypes derived from a single genome are shared between two distinct cell types.

**Summary:** The opportunistic human fungal pathogen *Candida albicans* switches between two distinct, heritable cell types, named “white” and “opaque.” We show that opaque cells, in response to proteins as the sole nitrogen source, up-regulate a specialized program, including specific secreted aspartyl proteases and peptide transporters. We demonstrate that, in mixed cultures, opaque cells enable white cells to respond and proliferate more efficiently under these conditions. These observations suggest that white-opaque switching creates mixtures of cells where the population characteristics - which derive from a single genome - reflect the contributions of two distinct cell types.

**Dataset Reference Numbers:** The .RAW files for both sets of Mass Spectrometry experiments have been deposited at the ProteoSAFe resource (https://proteomics.ucsd.edu/ProteoSAFe/).

MSP-MS experiment reference number: MSV000085279. For reviewer access use login “MSV000085279_reviewer” and password “candidamspms”.

Proteomics experiment reference number: MSV000085283. For reviewer access use login “MSV000085283_reviewer” and password “candidaprot”.

## Introduction

Protease secretion by pathogens plays an important role in many aspects of host-pathogen interactions, including colonization, tissue damage, and interference with host immune responses (Blom *et al.* 2009; Koziel and Potempa 2013; Pietrocola *et al.* 2017). Secreted proteases also contribute to nutrient acquisition; examples include amino acid acquisition by *Legionella pneumophila* (White *et al.* 2018) and carbon acquisition by *Pseudomonas aeruginosa* (Diggle *et al.* 2007). Secreted proteases are also important for the most prevalent fungal pathogen of humans, *Candida albicans*. The secreted aspartyl protease (*SAP*) family from *C. albicans* comprises thirteen homologous proteins and has been linked to, among other things, biofilm formation (Winter *et al.* 2016), interactions with bacteria (Dutton *et al.* 2016), adhesion to host cells (Watts *et al.* 1998; Bektic *et al.* 2001; Albrecht *et al.* 2006), protection from host defense proteins (Borg-von Zepelin *et al.* 1998; Gropp *et al.* 2009; Meiller *et al.* 2009; Rapala-Kozik *et al.* 2010, 2015; Bochenska *et al.* 2015; Kozik *et al.* 2015), and activation of host immune responses (Schaller *et al.* 2005; Hornbach *et al.* 2009; Pietrella *et al.* 2010; Pericolini *et al.* 2015; Gabrielli *et al.* 2016). The *SAP* family, especially *SAP2* (Hube *et al.* 1997), has also been linked to *C. albicans’* unusual ability, compared with other fungal species, to utilize proteins (e.g. casein or bovine serum albumin (BSA)) as a nitrogen source (Staib 1965; Nelson and Young 1986). This occurs through the cleavage of proteins into short peptides which are then imported by the oligopeptide transporter (*OPT*) and peptide transporter (*PTR*) families (Reuss and Morschhäuser 2006; Dunkel *et al.* 2013).

Although *C. albicans* is a common component of the human microbiome - asymptomatically colonizing the skin, gastrointestinal tract, and genitourinary tract of healthy individuals - it can cause superficial mucosal or dermal infections as well as disseminated bloodstream infections if the host immune system is compromised or the native microbiome is disrupted (Kennedy and Volz 1985; Wey *et al.* 1988; Wenzel 1995; Calderone and Fonzi 2001; Kullberg and Oude Lashof 2002; Eggimann *et al.* 2003; Gudlaugsson *et al.* 2003; Pappas *et al.* 2004; Achkar and Fries 2010; Kumamoto 2011; Kim and Sudbery 2011). *C. albicans* grows as several distinct cell types *in vitro* and *in vivo*, including yeast, pseudohyphal, and hyphal cells. In addition, *C. albicans* (and the closely related species *Candida dubliniensis* and *Candida tropicalis*) can switch between two distinct, heritable cell types named “white” and “opaque” (Slutsky *et al.* 1987; Soll *et al.* 1993; Johnson 2003; Pujol *et al.* 2004; Lohse and Johnson 2009; Soll 2009; Morschhäuser 2010; Porman *et al.* 2011). The white and opaque cell types are heritable for many generations, and switching between them occurs approximately once every 10,000 cell divisions under standard laboratory conditions (Rikkerink *et al.* 1988; Bergen *et al.* 1990). Although genetically identical, these two cell types differ in the appearance of their colonies, in their cell morphologies (Figure 1A), as well as in expression of roughly 15% of the genome (Lan *et al.* 2002; Tuch *et al.* 2010). As a result of these expression differences, the two cell types also differ in their abilities to mate (Miller and Johnson 2002), their responses to environmental conditions (Si *et al.* 2013), their metabolic preferences (Lan *et al.* 2002; Ene *et al.* 2016; Dalal *et al.* 2019), and their interactions with the host immune system (Kvaal *et al.* 1997, 1999; Geiger *et al.* 2004; Lohse and Johnson 2008; Sasse *et al.* 2013; Takagi *et al.* 2019). Transcriptional profiling has revealed that several *SAP*s and peptide transporters are differentially expressed between white and opaque cells (e.g. *OPT1* is enriched in white cells, *SAP1* and *OPT4* are enriched in opaque cells) (Hube *et al.* 1994; White and Agabian 1995; Lan *et al.* 2002; Tuch *et al.* 2010; Hernday *et al.* 2013). It has also been shown that opaque cells proliferate better than white cells when di-peptides, tri-peptides, or BSA are the sole nitrogen source (Kvaal *et al.* 1999; Lan *et al.* 2002; Ene *et al.* 2016).

**Figure 1:**
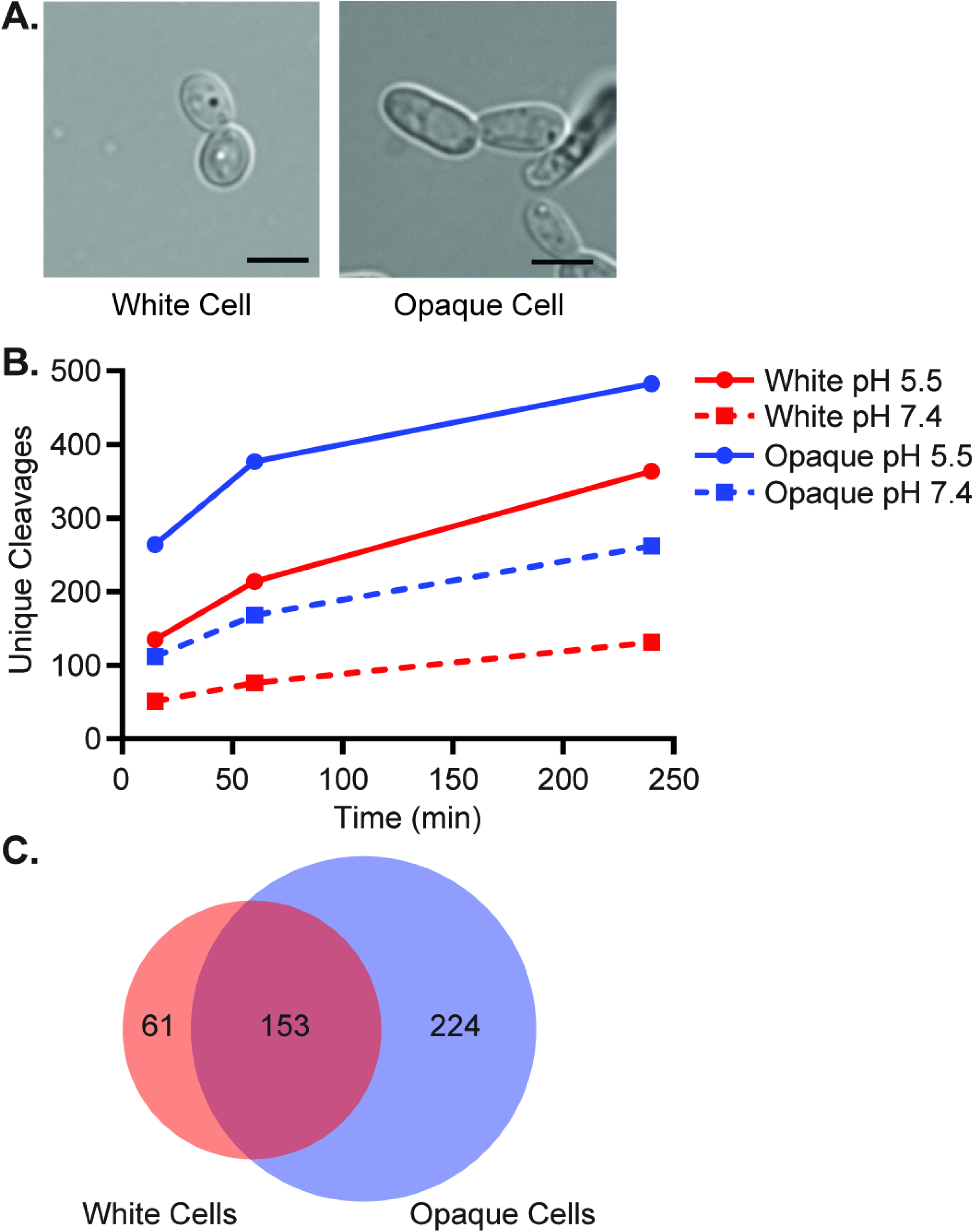
Opaque cells secrete higher and more diverse proteolytic activity than white cells. (A) Images of typical white (left) and opaque (right) *C. albicans* cells grown in liquid SCD+aa+Uri media at 25°C. Scale bar is 5 µm, panel adapted from Lohse and Johnson 2016 (Lohse and Johnson 2016). (B) Comparison of the number of unique 8-mer cleavages seen in white (red) and opaque (blue) cell conditioned media at pH 5.5 (solid lines, circles) and pH 7.4 (dashed lines, squares) after 15, 60, and 240 minutes. (C) Quantification of the number of unique 8-mer cleavages common to both cell types or specific to one cell type (pH 5.5, 60 minutes). All MSP-MS cleavages are compiled in File S3.

In this paper, we show that opaque cells have a specialized program that is induced when protein present in the environment is the sole source of nitrogen. This inducible program, which is not observed in white cells, includes the massive up-regulation of several specific SAP proteins along with a series of peptide transporters. We show that this induced response is needed for the ability of opaque cells to efficiently use proteins as a sole nitrogen source and, using a series of quadruple and quintuple deletion mutants, we identify the individual SAPs needed for opaque cells to proliferate under these conditions. We also show that Stp1, a known regulator of *SAP2* in white cells (Martínez and Ljungdahl 2005), works in combination with the white-opaque regulators Wor3 and Efg1 to produce the specialized opaque proteolysis program. Finally, we show that, in mixed cultures of white and opaque cells, opaque cells, even when in the minority, can enable white cells to proliferate efficiently when proteins are the sole nitrogen source. Based on these results, we suggest that one role of white-opaque switching is to create mixed populations where the population characteristics derived from a single genome are shared between two distinct cell types.

## Materials and Methods

Supplemental Materials and Methods, Supplemental Results, as well as legends for all Supplemental Figures, Files, and Tables can be found in File S1.

### Media and Growth Conditions

Unless otherwise noted, strains were grown at 25°C on synthetic complete media (6.7 g/L Yeast Nitrogen Base without Amino Acids (BD #291940)) supplemented with 2% glucose, amino acids (2 g/L), and uridine (100 µg/mL) (SCD+aa+Uri); plates contained 2% Agar. Specific nitrogen source and nitrogen depletion media was prepared immediately before use from 2X S+Uri (3.4 g/L Yeast Nitrogen Base without Amino Acids or ammonium sulfate (MP Biomedicals, #4027-032), 200 µg/mL uridine), 40% Glucose, 5% ammonium sulfate (Sigma A2939), and 10% protein (BSA (Sigma A1470), HSA (Sigma A8763), Hemoglobin (Sigma H2500), or Myoglobin (Sigma M1882)) stocks. Final glucose, ammonium sulfate, and protein concentrations, when present, were 2%, 0.5%, and 1% respectively. Common combinations included BSA without ammonium sulfate or amino acids (SD+BSA+Uri), ammonium sulfate without BSA or amino acids (SD+AmS+Uri), or media lacking BSA, ammonium sulfate, and amino acids (SD+Uri). Specific nitrogen source, nitrogen depletion, and specific carbon source liquid media were filtered on 0.45 µm PES filters (Thermo #725-2545) prior to use. Dulbecco’s PBS (D-PBS) lacking calcium chloride and magnesium chloride was procured from the Cell Culture facility at the University of California, San Francisco or Gibco (#14190-136). A list of media used in this study can be found in File S2.

### Plasmid Construction

The mCherry reporter plasmids were constructed as follows in the pUC19 backbone. In three separate steps, *C. albicans* optimized mCherry taken from pMBL180 (Lohse and Johnson 2016), the *SAT1* selectable marker from pNIM1 (Park and Morschhäuser 2005), and the *RPS1*/*RP10* homology region from pADH33 (Lohse *et al.* 2013) were inserted between the HindIII-PstI, PstI-BamHI, and BamHI-EcoRI restriction sites, respectively, to create the plasmid pNZ116. The promoters for *OPT1* (1968bp), *OPT2* (2056bp), *SAP2* (4522bp), and *UGA4* (980bp) were amplified from *C. albicans* SC5314 genomic DNA and added to HindIII digested pNZ116 backbone using In-Fusion cloning (Takara #638911) to generate plasmids pNZ119 (*OPT1*), pNZ120 (*OPT2*), pNZ121 (*SAP2*), and pNZ118 (*UGA4*). The promoter regions extend from the start or stop of the upstream gene (with the exception of p*OPT2* and p*UGA4*, which start 24 and 6 base pairs downstream of that location, respectively) to the ATG at the start of the respective genes. All plasmid sequences were verified by sequencing. A list of oligonucleotides and plasmids used in this study can be found in File S2.

### Strain Construction

The SC5314-derived *C. albicans* wild type white and opaque strains used in the proteomic and MSP-MS experiments have been previously reported (Hernday *et al.* 2013), in brief these are *HIS1* and *LEU2* addbacks to the SN152 **a**/α *his1 leu2 arg4* strain (Noble and Johnson 2005) that were then converted to the switching capable **a**/Δ background by deletion of the α copy of the Mating Type Like (*MTL*) locus using pJD1 (Lin *et al.* 2013). The opaque transcriptional regulator knockout library strains (Lohse *et al.* 2016) derive from the same background. The previously reported white and opaque Tef2-GFP and Tef2-mCherry strains (Takagi *et al.* 2019) are derived from the switching capable AHY135 strain (Hernday *et al.* 2013), where the *HIS1* and *LEU2* markers were added back to RZY47, itself a sorbose selected **a**/**a** copy of the SN87 **a**/α *his1 leu2* strain (Noble and Johnson 2005; Zordan *et al.* 2006). *C. albicans* clinical isolates L26 and P37005 (Lockhart *et al.* 2002), *C. dubliniensis* CD36 (Sullivan *et al.* 1995), *C. tropicalis* MYA3404 (Joly *et al.* 1996), the *C. tropicalis* AM2005/0093 derived white-opaque switching strains (Anderson *et al.* 2016), and *Candida parapsilosis* CBS604/ATCC22019 (Guerin *et al.* 1989) have all been previously reported.

Gene deletion and truncation utilized the *SAT1* marker-based CRISPR protocol targeting *Candida maltosa LEU2* described by Nguyen and colleagues (Nguyen *et al.* 2017). We used a derivative of the hemizygous *LEU2* strain SNY250 (a derivative of SNY152 (**a**/α *his1 leu2 arg4*) with the *C. dubliniensis HIS1* and *C. maltosa LEU2* gene deletion cassettes integrated at the *C. albicans LEU2* locus) (Noble and Johnson 2005) which was converted to **a**/Δ by deleting the α copy of the MTL using pJD1 (Lin *et al.* 2013). For gene deletions, the 90bp-annealed donor DNA (dDNA) contains homology to the regions directly upstream and downstream of the targeted ORF. Each dDNA homology arm consists of 44 bp and the two homology arms are separated by a two base pair GG insert added to create a potential gRNA site. For the *STP1* truncation mutant (Δ2-61), the dDNA homology arms flank the targeted amino acids and no GG insert was added. Gene deletions were confirmed by colony PCR reactions verifying the loss of the targeted ORF(s) and truncation mutations were confirmed by sequencing. After confirming the presence of the desired edit(s), the Cas9 ORF-gRNA-*SAT1* cassette was recycled by plating on Leu/His/Arg dropout plates and selecting for recombination events with an intact *CmLEU2* ORF. We selected against both leucine and histidine in order to avoid potential histidine auxotrophies arising during the recombination process (both *CmLEU2* and *CdHIS1* are present at the *CaLEU2* locus in the SNY250-derived background).

The mCherry reporter strains were constructed using AgeI-HF (NEB R3552L) linearized reporter plasmids transformed into the SNY250-derived **a**/Δ wild type strain. Colonies were selected for growth on Yeast Extract Peptone Dextrose (YEPD) plates supplemented with 400 μg/mL nourseothricin (clonNAT, WERNER BioAgents, Jena, Germany). Plasmid integration at the *RPS1* (*RP10*) locus was verified by colony PCR across the 5’ and 3’ flanks of the integrated plasmid.

A list of oligonucleotides and strains used in this study can be found in File S2.

### Conditioned Media Processing for MSP-MS and Proteomic Analyses

Following recovery from glycerol stocks, white and opaque *C. albicans* strains were grown for seven days on SCD+aa+Uri plates at 25°C. Overnight cultures (5 mL, SCD+aa+Uri, 25°C) were started from single colonies with no visible switching events. The following morning, cell type homogeneity of the overnight culture was verified by microscopy prior to dilution to OD_600_ = 0.05 in 50 mL of SCD+aa+Uri media in a 250 mL flask; two independent 50 mL cultures were grown for each strain. Cultures were incubated with shaking at 220 rpm for 24 hours at 25°C before harvesting as previously described for planktonic *C. albicans* cultures (Winter *et al.* 2016). In short, cultures were transferred to 50 mL tubes and centrifuged for 10 minutes at 3500rpm at 4°C. The supernatant was collected, filtered on a 0.45 µm PES filter (Thermo #725-2545), and flash frozen in liquid nitrogen. A small aliquot was taken from each culture immediately prior to harvesting, diluted, and plated on three SCD+aa+Uri plates which were incubated for seven days at 25°C and then scored for colony morphology to confirm each culture’s cell type.

To prepare samples for analysis, the frozen conditioned media was thawed on ice, the conditioned media from the two independent cultures of each strain were pooled, and the combined sample was concentrated (using a refrigerated centrifuge) to approximately 1 mL on 10kDa MWCO Amicon Ultra spin filter units (Millipore UFC901024). The concentrated solutions were then diluted to 15 mL with ice cold D-PBS to exchange the buffer and subsequently concentrated to approximately 750 µL. These concentrated solutions were then aliquoted, flash frozen in liquid nitrogen, and stored at −80°C. Protein concentrations of each solution were quantified using the Bradford assay.

### MSP-MS Analysis

Substrate specificity profiles were determined for 24 hour conditioned media samples, prepared as described above, from white and opaque cultures from an SC5314-derived *C. albicans* strain using the previously reported MSP-MS assay (O’Donoghue *et al.* 2012; Winter *et al.* 2016, 2017). In short, 20 µg/mL processed conditioned medium and matched no-enzyme controls were assayed at room temperature (approximately 22°C) against a diverse library of 228 tetradecapeptides pooled at 500 nM in D-PBS (pH 7.4) and MES (pH 5.5; 9.5 mM MES, 2.7 mM KCl, 140 mM NaCl). 30 µL of each 150 µL assay mixture was removed after 15, 60, and 240 minutes, quenched with 30 µL quanidinium-HCl, and flash frozen in liquid nitrogen. Samples were thawed, acidified to pH 2 by the addition of 1.5 µL of 20% formic acid, desalted with C_18_ Desalting Tips (Rainin 17014047), eluted in 40 µL of a 50:50 acetonitrile (Sigma 34851) : water (Fisher W5) mixture with 0.2% formic acid, and then lyophilized. Samples were resuspended in 30 µL of 0.2% formic acid solution prior to mass spectrometry.

Cleavage site identification was performed on an LTQ Orbitrap XL mass spectrometer (Thermo) equipped with a nanoACQUITY (Waters) ultraperformance LC (UPLC) system and an EASY Spray ion source (Thermo). Reversed-phase chromatography was carried out with an EASY-Spray PepMap C_18_ column (Thermo, ES800; 3 µm bead size, 75 µm by 150 mm). Loading was performed at a 600 nL/min flow rate for 12 min, and then peptide separation was performed at a 300 nL/min flow rate over 63 min with a linear gradient of 2 to 30% (vol/vol) acetonitrile in 0.1% formic acid followed by a 2 min linear gradient from 30 to 50% acetonitrile. Peptide fragmentation was performed by collision-induced dissociation (CID) on the six most intense precursor ions, with a minimum of 1,000 counts, with an isolation width of 2.0 *m/z* and a minimum normalized collision energy of 25. For MS/MS analysis, survey scans were recorded over a range of 325 to 1,500 *m/z*. Internal recalibration to polydimethylcyclosiloxane ion (*m/z* =445.120025) was used for both MS and MS/MS scans. MS peak lists were generated with MSConvert. Data were searched against the 228-member peptide library using the Protein Prospector software (http://prospector.ucsf.edu/prospector/mshome.htm, UCSF) (Chalkley *et al.* 2008) with specified tolerances of 20 ppm for parent ions and 0.8 Da for fragment ions. All cleavages were allowed in the search by designating “no enzyme” specificity. The following variable modifications were used: amino acid (proline, tryptophan, and tyrosine) oxidation and N-terminal pyroglutamate conversion from glutamine. Protein Prospector score thresholds were selected with a minimum protein score of 15 and a minimum peptide score of 15. Maximum expectation values of 0.01 and 0.05 were selected for protein and peptide matches, respectively. Peptides corresponding to cleavage products in the 228-member library (O’Donoghue *et al.* 2015) were selected with in-house software and imported into iceLogo software v.1.2 (Colaert *et al.* 2009) to generate substrate specificity profiles and Z score calculations as described previously (O’Donoghue *et al.* 2012). Octapeptides corresponding to P4-P4’ were used as the positive data set, and octapeptides corresponding to all possible cleavages in the library (*n*=2,964) were used as the negative data set. Octapeptide cleavage products identified by the MSP-MS assays with the 228-member library are provided in File S3 in the supplemental material.

### Proteomic Analysis

Proteomic analysis of conditioned media from planktonic cultures of white and opaque cells from three *C. albicans* backgrounds (SC5314, L26, P37005) was based on previously reported methods (Winter *et al.* 2016). In brief, three samples from 24 hour conditioned media preparations of each strain (3 µg diluted to 40 µL with D-PBS) were prepared as described above and matched on a total protein basis. Conditioned media preparations were incubated for 20 minutes with 6 M urea (Sigma U5378) and 10 mM DTT (Sigma D0632) at 55°C, alkylated for 1 hr with 12.5 mM iodoacetamide (Sigma I6125) at room temperature (approximately 22°C), quenched with an additional 10 mM DTT, and diluted 2.5-fold with 25 mM ammonium bicarbonate (100 µL, Sigma A6161). For trypsinization, 16 µL of a 1:100 dilution (in 25 mM ammonium bicarbonate) of trypsin solution (Promega V511C) was added, and the digests were incubated for 17 hours at 37°C (1:37.5 trypsin/sample w/w). Following the trypsin digest, samples were acidified to pH 2 with 5 µL of 20% formic acid (JT Baker 0128-01) before desalting with C_18_ Desalting Tips (Rainin 17014047). Samples were eluted in 40µL of a 50:50 acetonitrile (Sigma 34851) : water (Fisher W5) mixture with 0.2% formic acid and then lyophilized. Samples were resuspended in 50 µL of 0.2% formic acid solution prior to mass spectrometry.

LC-MS/MS peptide sequencing was performed using the LTQ Orbitrap-XL mass spectrometer, ion source, UPLC system, EASY-Spray PepMap C_18_ column, and CID parameters for peptide fragmentation described above. Loading was performed at a 600 nL/min flow rate for 12 min, and then peptide separation was performed at a 300 nL/min flow rate over 63 min with a linear gradient of 2 to 30% (vol/vol) acetonitrile in 0.1% formic acid followed by a 2 min linear gradient from 30 to 50% acetonitrile. For MS/MS analysis, survey scans were recorded over a mass range of 325 to 1,500 *m/z*. MS peak lists were generated with in-house software called PAVA. Database searching was performed with Protein Prospector against the UniProtKB *C. albicans* database [SC5314, taxonomic identifier 237561; downloaded 20 April 2017 (last modified 4 February 2017) with 6,153 entries]. The databases were concatenated with an equal number of fully randomized entries for estimation of the false-discovery rate (FDR). Database searching was carried out with tolerances of 20 ppm for parent ions and 0.8 Da for fragment ions. Peptide sequences were matched as tryptic peptides with up to two missed cleavages. Constant and variable modifications were set as described previously (O’Donoghue *et al.* 2012). The following Protein Prospector score thresholds were selected to yield a maximum protein FDR of less than 1%. A minimum “protein score” of 22 and a minimum “peptide score” of 15 were used; maximum expectation values of 0.01 for protein and 0.05 for peptide matches were used; Report Homologous Proteins was set to “Interesting”. Only proteins with a minimum of two unique peptides for identification are reported.

Each protein’s individual peptide counts from the three samples for a given strain were averaged, using a value of zero when no peptides were detected in a given sample. The percent abundance of a protein in a strain was calculated as the average spectral count from the three samples divided by the sum of the average spectral counts for all proteins detected in that strain. A complete list of proteins identified by these experiments are provided in File S4 in the supplemental material. We note that three proteins (Bgl2, Fet31, Rny11) appeared twice in the protein list for at least one sample; we have left both instances in place in File S4 and for the percent abundance calculations. Omitting the second entries of these proteins had less than a 0.1% effect on the abundance of all SAPs expect for the most highly expressed samples (Sap1, Sap3, Sap98, Sap99 in opaque cells) where the largest effect was 0.66%; none of our conclusions are affected by the inclusion and/or exclusion of these entries.

### Conditioned Media BSA Cleavage Assays

In order to examine basal proteolytic activity, 3 mL SCD+aa+Uri cultures were started from white or opaque colonies and incubated for 24 hours at 25°C on a roller drum. The pH of SCD+aa+Uri media, as evaluated by pH paper (Baker #4393-01 and #4391-01), decreased from approximately 5.0-5.5 to 3.0-3.5 during the 24 hour incubation, we note that this range is compatible with the reported pH preferences of much of the SAP family (Koelsch *et al.* 2000; Aoki *et al.* 2011). After the 24 hour incubation, supernatant was collected and filtered on a 0.45 µm PES filter (Thermo #725-2545). BSA was added to the filtered supernatant at a final concentration of 0.5% (from a 20% stock solution) and the mixture was vortexed and incubated for two hours at 25°C in a glass test tube on a roller drum. Samples were collected immediately after the addition of BSA and after 15, 60, and 120 minutes. The test tubes were vortexed prior to harvesting, and the harvested samples were mixed 1:4 with 8 M Urea, vortexed, and flash frozen in liquid nitrogen. Samples were run on 12.5% SDS-PAGE gels to evaluate effects on the full length BSA band. When indicated, Pepstatin A (Sigma P5318) was diluted from 5 mM (in DMSO) to 1 mM (in DMSO) before being added to the filtered conditioned media (prior to addition of BSA) at a final concentration of 1 or 5 µM as indicated, equivalent volumes of DMSO were used as loading controls for these assays.

When screening the opaque transcriptional regulator knockout library, we processed 16 strains at a time and only harvested samples after 120 minutes. Candidates with reduced cleavage of BSA identified by the initial screen were subjected to further testing in order to control for experimental artifacts such as predominately white cell populations or lower culture densities. First, we repeated the assay while microscopically verifying the cell type of each culture. This eliminated the white cell population concern for all but two strains for which we could not get majority opaque populations (*bcr1, fgr15*). We then repeated the assay with the remaining candidates and, after verifying cell type, determined the cell density of each sample by flow cytometry. To compensate for the lower cell density observed for several strains after 24 hours, the assay was repeated with cultures incubated for 40 or 44 hours. After accounting for cell type and culture density, only the *efg1* and *wor3* deletions exhibited a BSA cleavage-deficient phenotype.

### Flow Cytometry Proliferation and Expression Assays

Flow Cytometry assays used either a BD Accuri C6 Plus (basic proliferation assays, GFP-based co-culture assays) or a BD FACS Celesta (mCherry-based co-culture and Proteinase K pretreatment reporter assays). The BD Accuri C6 Plus used a 488nm laser with a 533/30 bandpass filter, samples were loaded from 96-well U-bottom plates (Falcon 351177), and normal acquisition was 10 μL of sample on the slow setting. The BD FACS Celesta used a 488nm laser with a 530/30 bandpass filter and a 561nm laser with a 610/20 bandpass filter, samples were loaded from 5 mL tubes (Falcon 352052) and normal acquisition was 10,000 events on the high setting.

For flow cytometry proliferation assays, SCD+aa+Uri overnight cultures (25°C) were started from white or opaque colonies of each strain; independent cultures were processed in parallel for repeats of a given strain or condition. The following morning, cultures were diluted back to OD_600_ = 0.5 in 5 or 15 mL fresh SCD+aa+uri media and grown for five to ten hours, depending on the assay, at 25°C in a roller drum. Cells were then harvested, washed twice with 4 mL D-PBS, resuspended in SD+Uri, and diluted to OD_600_ = 0.5 in SD+Uri. Cell densities were verified by flow cytometry and the cells were then diluted 1:10 to OD_600_ = 0.05 in the appropriate media (e.g. SD+BSA+Uri or SD+AmS+Uri) and gently vortexed. The density of each culture was then determined by flow cytometry and the strains were grown on a roller drum at 25°C. At each subsequent time point, cultures were gently vortexed, samples were removed, and denser samples were diluted with D-PBS prior to determining cell density by flow cytometry. Cell counts for proliferation assays were based on the number of cells detected in a 10 μL sample multiplied by the sample’s dilution. For the assay comparing proliferation on different proteins (BSA, HSA, Hemoglobin, Myoglobin), the initial 10% protein stocks were dialyzed at 4°C in 3,500 MWCO Slide-A-Lyzer Dialysis Cassettes (Thermo #66330) versus distilled water for approximately eight hours (roughly 1:250) and then against fresh distilled water for a further sixteen hours (again, roughly 1:250) prior to incorporation into media. For the 37°C BSA proliferation assay, samples were incubated in a 37°C, rather than a 25°C, roller drum and cells were harvested for plating for cell type determination as described in File S1. For the BSA as a carbon source assays, cells were diluted to OD_600_ = 0.5 in S+Uri and then 1:10 to OD_600_ = 0.05 in either S+BSA+Uri or S+BSA+AmS+Uri media.

#### Flow Cytometry Co-Culture Proliferation Assays

The co-culture proliferation assays followed the protocol described above with the following modifications. After the D-PBS washes, SD+Uri resuspension, and OD_600_ determination, the two strains were mixed at defined Tagged / Untagged ratios (e.g. 100/0, 50/50, 20/80, 5/95, and 0/100 for the wild type opaque and white experiments) where the two strains had a combined starting OD_600_ of 0.5. Expression of a Tef2-GFP fusion protein was used to distinguish strains, unless otherwise noted this reporter was expressed by the wild type opaque strain.

#### Flow Cytometry mCherry Reporter Assays

The co-culture mCherry reporter proliferation assays followed the protocol described above with the following modifications. After the D-PBS washes and SD+Uri resuspension, the initial dilution in SD+Uri was to OD_600_ = 1.0, rather than 0.5, and the subsequent 1:10 dilution into the appropriate media was to OD_600_ = 0.1, rather than 0.05. Cells were analyzed using a BD FACS Celesta rather than a BD Accuri C6 Plus. The Tef2-GFP fusion protein was used to distinguish between the opaque cells (GFP positive) and the white strains expressing the mCherry reporters for *OPT1, OPT2, SAP2*, or *UGA4* expression (GFP negative). For the Proteinase K pretreatment assays, 10 mL of a 10% BSA solution was incubated with 2.5 μL of Proteinase K (NEB P8107S) for 30 minutes at room temperature (∼22°C). The Proteinase K-treated BSA, in parallel with a mock treated BSA control, was then used to make SD+BSA+Uri media as described above.

#### Flow Cytometry Analysis

Data for each sample was exported as an FCS file, all subsequent analysis was done with R (R Core Team 2019) using the flowCore (Ellis *et al.* 2019), ggcyto (Van *et al.* 2018), and tidyverse (Wickham 2017) packages. For the myoglobin samples, which accumulated debris during the experiment, the data were filtered during analysis to exclude events with a FSC of less than 200,000. For the co-culture experiments performed on the Accuri (Figures 6B, S12, S15), data were first filtered to exclude those with no GFP fluorescence measurement. Cells were defined as GFP positive if the ratio of GFP.A/SSC.A was greater than 0.04. For the co-culture and/or reporter experiments performed on the Celesta (Figures 6C, S13, S14), cells were defined as GFP positive if the ratio of GFP.A/SSC.A was greater than 0.5. In both sets of co-culture analysis, the GFP positive evaluation was performed on all samples (including those that should contain 100% or 0% GFP positive cells).

**Figure 2:**
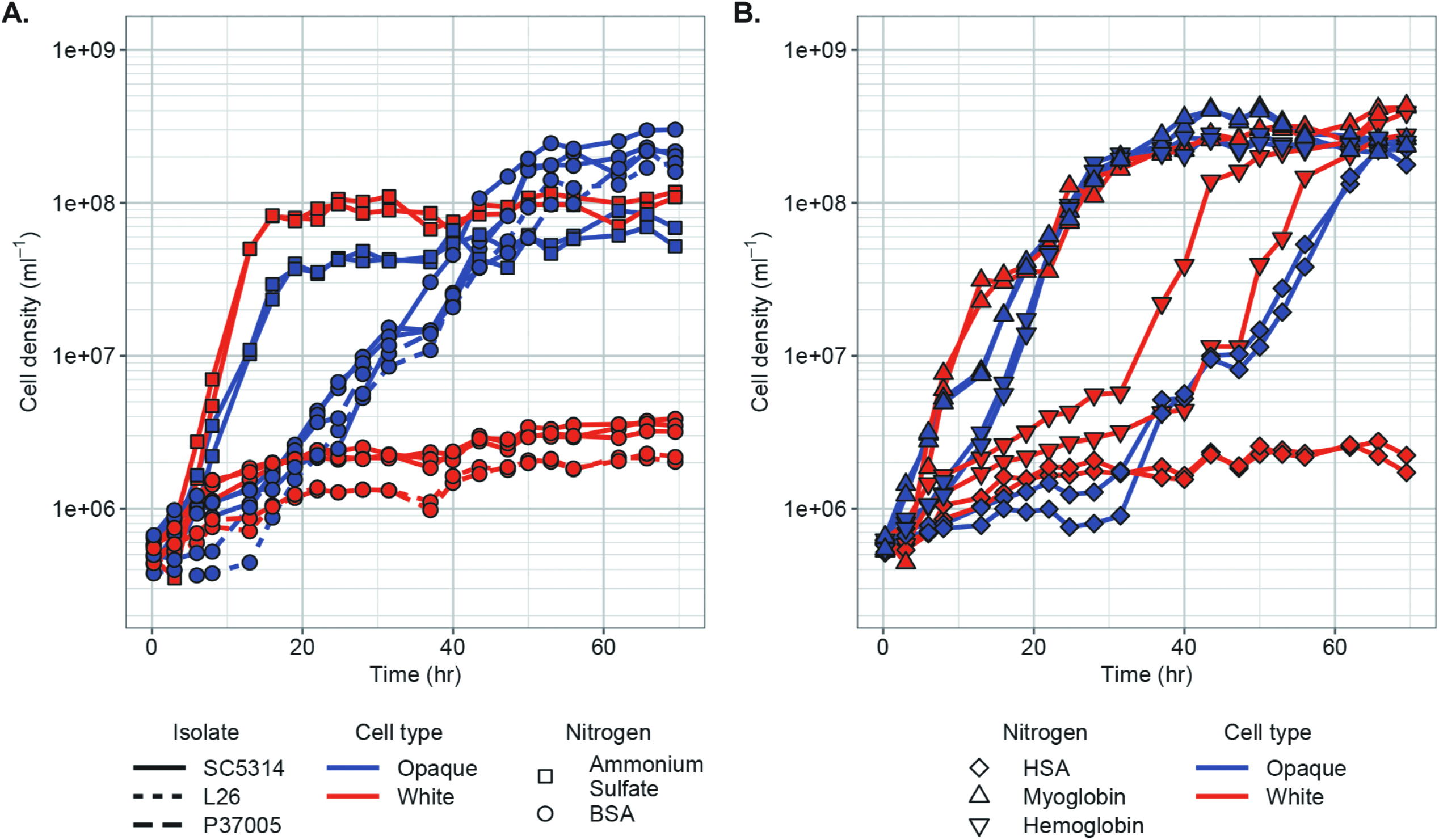
Opaque cells are more efficient than white cells at utilizing protein as the sole nitrogen source. (A) Proliferation of white (red) and opaque (blue) cells from three backgrounds (SC5314, solid line; L26, short dash; P37005, long dash) when ammonium sulfate (squares) or BSA (circles) is the sole nitrogen source. (B) Proliferation of white (red) and opaque (blue) cells from the SC5314 background when HSA (diamonds), myoglobin (upward triangles), or hemoglobin (downward triangles) is the sole nitrogen source. Cell counts in both panels were determined by flow cytometry.

**Figure 3:**
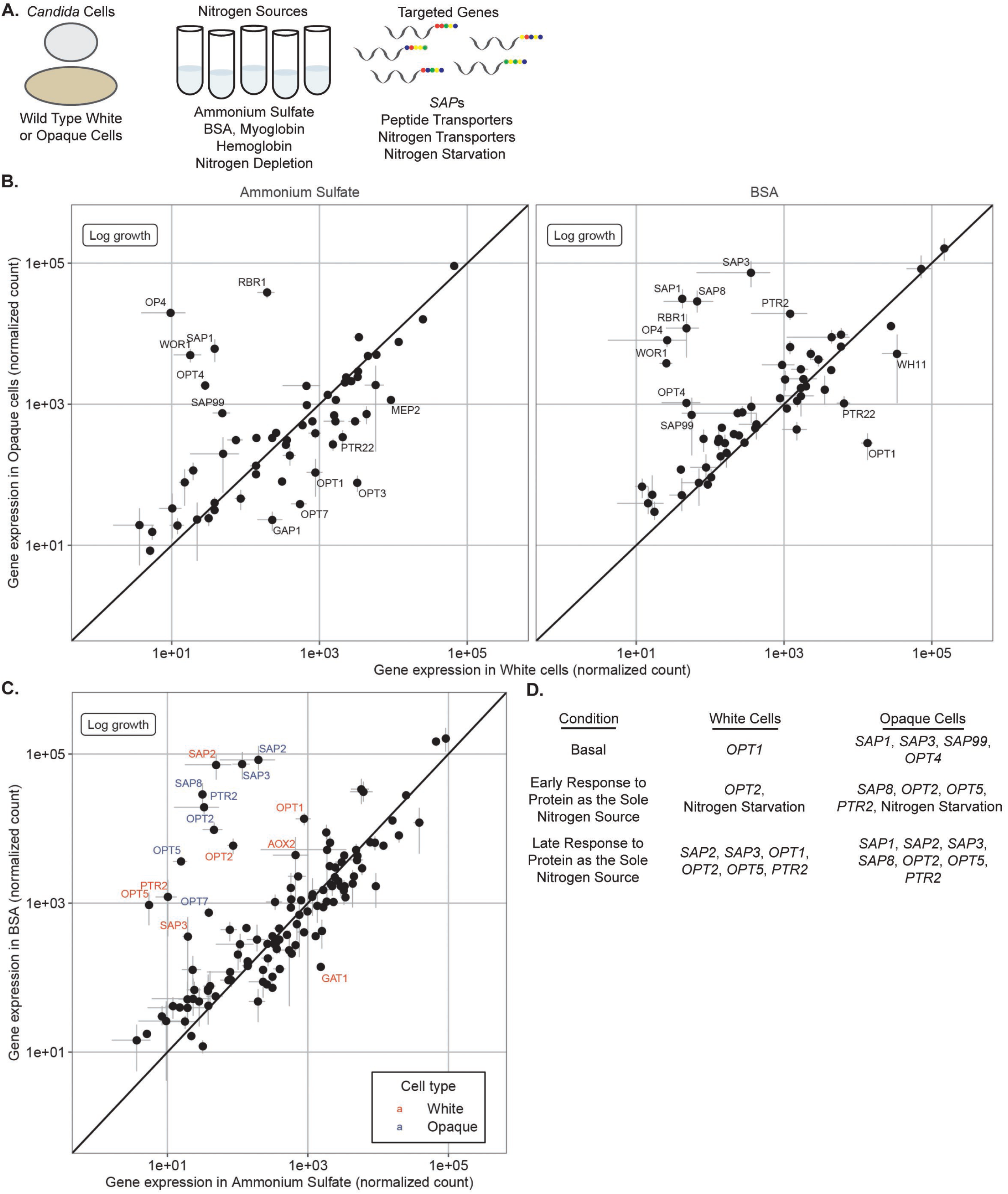
White and opaque cells express different *SAP*s and peptide transporters in response to different nitrogen sources. (A) NanoString probes were used to measure expression of the *SAP* family, transporters, and genes associated with nitrogen starvation. (B) White (lower right) and opaque (upper left) enrichment of selected genes in cells undergoing logarithmic proliferation on ammonium sulfate (left chart) or BSA (right chart) as the sole nitrogen source. Names are indicated for genes differentially regulated at least 6-fold between cell types, the line (y = x) indicates equal expression in both cell types. (C) Enrichment of selected genes in cells undergoing logarithmic proliferation on ammonium sulfate (lower right) or BSA (upper left) as the sole nitrogen source. Names are indicated for genes differentially regulated at least 6-fold between the two nitrogen sources in white (red) and opaque (blue) cells, the line (y = x) indicates equal expression on both nitrogen sources. (D) Summary of *SAP*s and transporters that are basally expressed and/or expressed when extracellular proteins are the sole nitrogen source.

**Figure 4:**
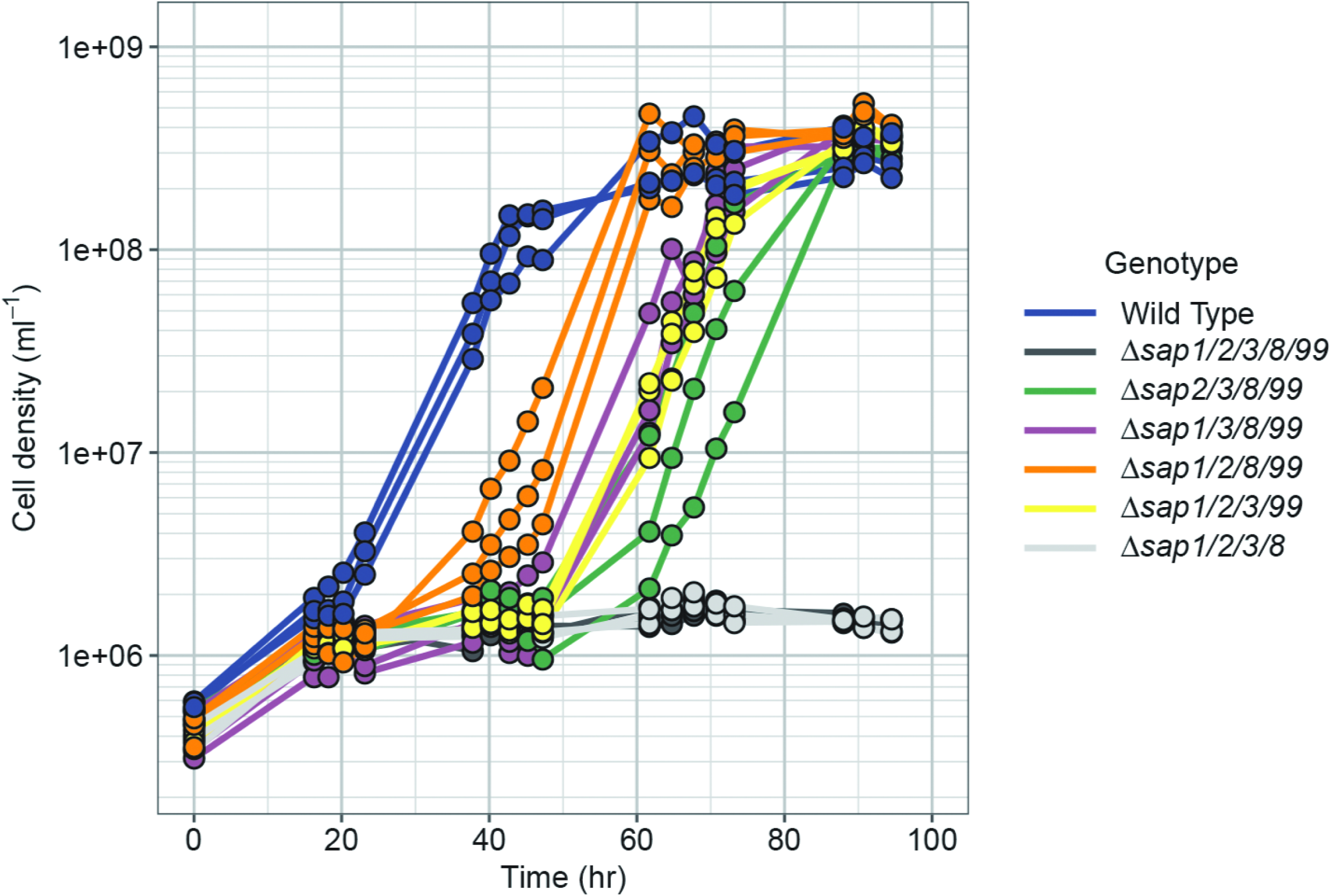
Any one of four *SAP* genes (*SAP1, SAP2, SAP3, SAP8*) is sufficient for opaque cell proliferation when BSA is the sole nitrogen source. Proliferation of opaque strains with combinations of either four or five of the five opaque-expressed *SAP*s (*SAP1, SAP2, SAP3, SAP8*, and *SAP99*) deleted on media where BSA is the sole nitrogen source. Cell counts were determined by flow cytometry.

**Figure 5:**
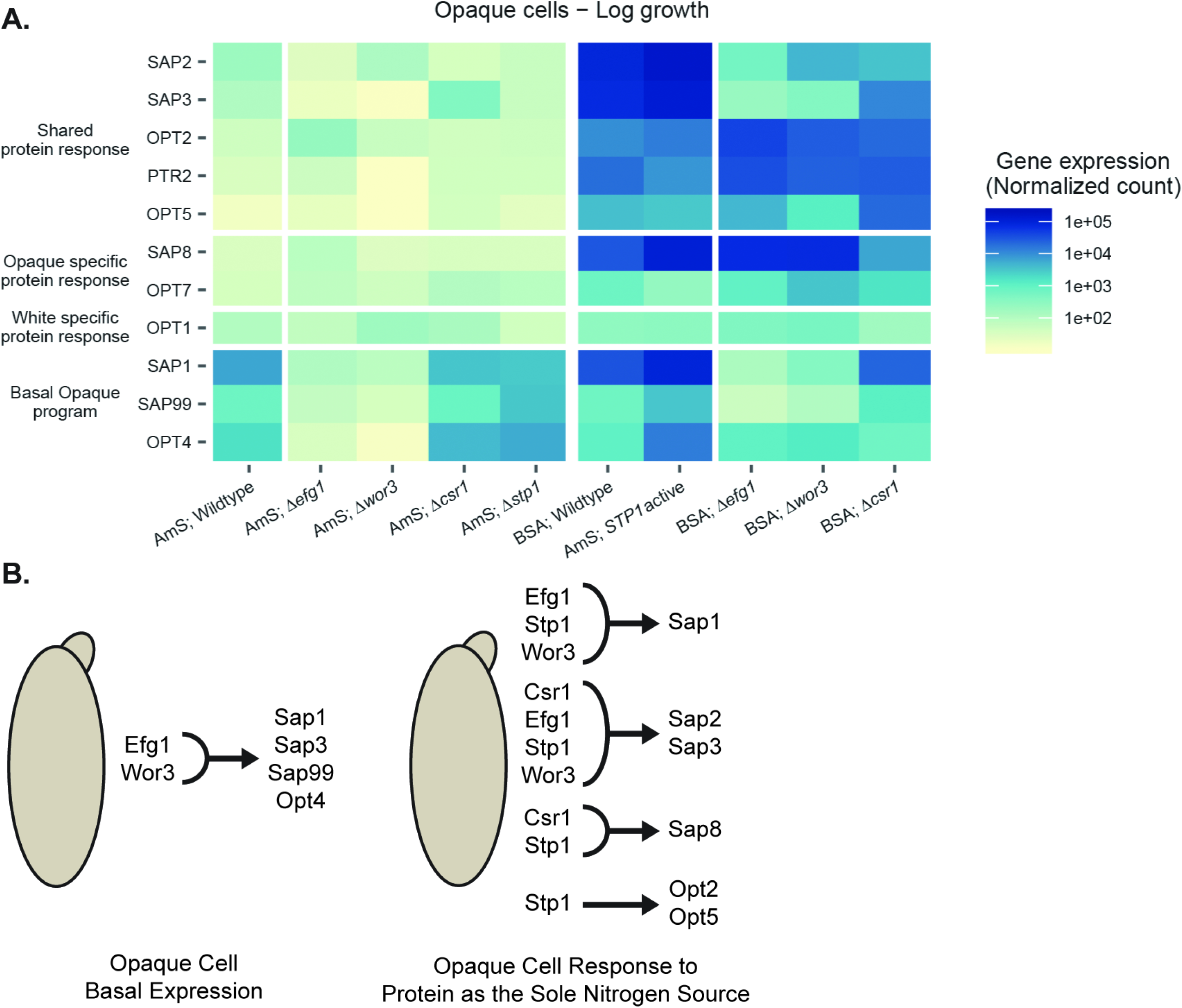
In opaque cells, four transcriptional regulators are necessary for either basal proteolytic activity, gene induction in response to proteins as a nitrogen source, or proliferation when proteins are the sole nitrogen source. (A) Transcript levels of selected genes in opaque cells of indicated genetic backgrounds growing on media where ammonium sulfate or protein (BSA) are the sole nitrogen sources. Efg1 and Wor3, but not Csr1 or Stp1, are necessary for the basal opaque cell *SAP* and transporter expression profile when ammonium sulfate is the sole nitrogen source. Wor3, Efg1, and Csr1 are needed for the full elaboration of the *SAP* component of the response to proteins in opaque cells. (B) Summary of the roles of Stp1, Efg1, Wor3, and Csr1 in the regulation of the basal opaque and opaque protein as sole nitrogen-source-induced expression of *SAP*s and transporters.

**Figure 6:**
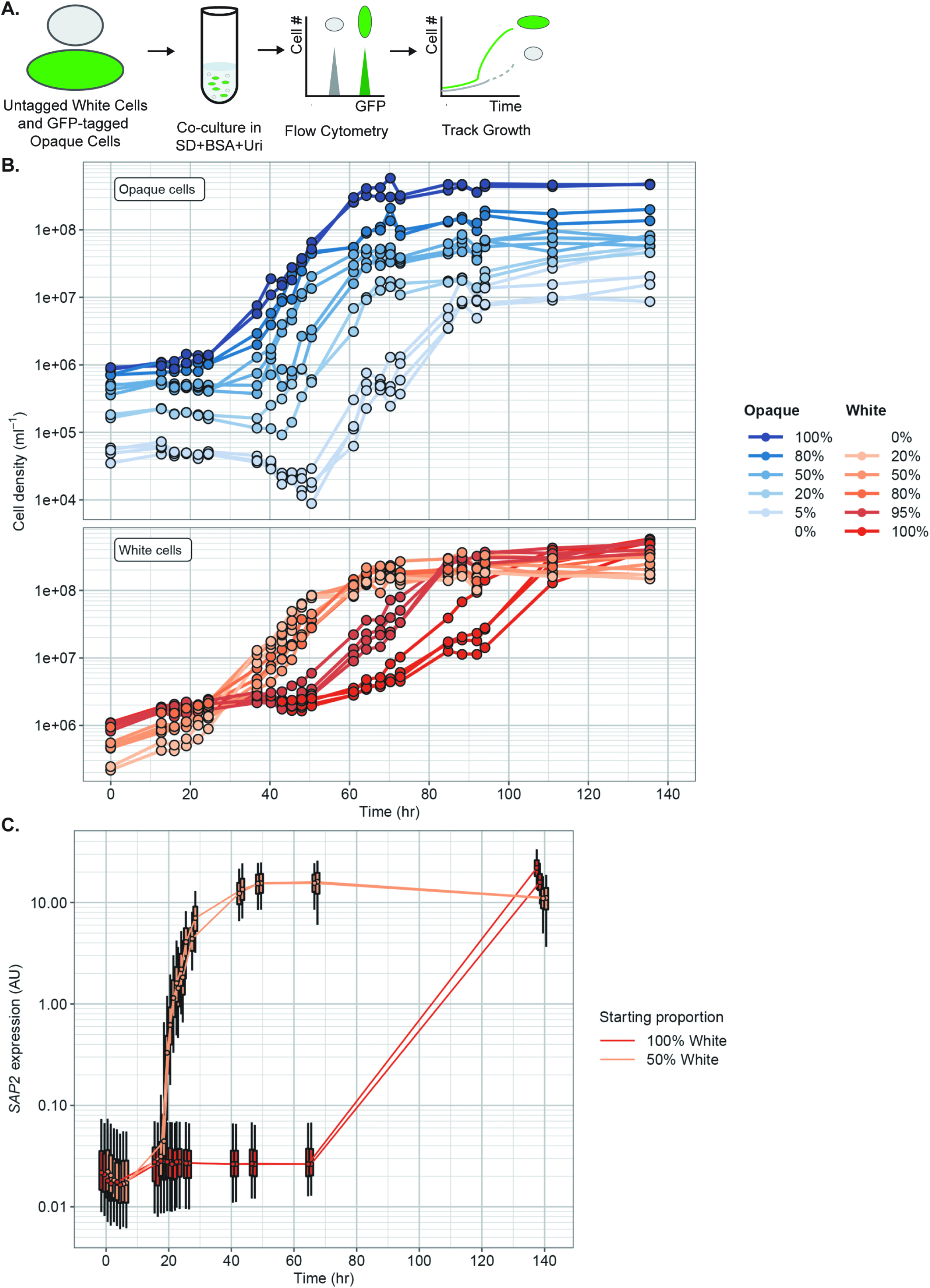
White cells more rapidly utilize BSA as a sole nitrogen source in the presence of opaque cells. (A) Different ratios of GFP-tagged opaque cells and untagged white cells were co-cultured in SD+BSA+Uri media at 25°C. Changes in cell density were measured by flow cytometry over the course of several days and the two cell types were distinguished by GFP fluorescence. (B) Proliferation of opaque (top, shades of blue) and white (bottom, shades of red) cells grown by themselves or co-cultured at different ratios when BSA is the sole nitrogen source. Cell counts and GFP fluorescence were determined by flow cytometry. (C) Expression of a mCherry reporter driven by the *SAP2* promoter in white cells, grown by themselves (red) or with an equal number of opaque cells (orange), when BSA is the sole nitrogen source. Boxes indicate the 25^th^ to 75^th^ percent of the data and whiskers indicate the 5^th^ to 95^th^ percent of the data for each sample at each time point. mCherry fluorescence was determined by flow cytometry and normalized by side scatter.

Growth rates were calculated using a sliding window approach; the natural log of cell count was regressed against time for each set of five time points based on previously reported protocols (Ziv *et al.* 2013). The growth rate was calculated as the greatest slope of a regression that had both an R^2^ greater than 0.9 and at least a four-fold change over the five time points. Subsequently, lag duration was estimated as the intersection of this regression with a horizontal line determined by the cell count at the first time point. Confidence intervals for mean growth rate and lag duration were calculated as +/- 1.96 the standard error of the mean. Growth rates calculated based on Flow Cytometry Proliferation Assays are included in File S5.

### White-Opaque Switching Assay and Variants

The white-to-opaque and opaque-to-white switching assays evaluating the effect of *SAP* deletions on switching rates followed previously reported protocols (Miller and Johnson 2002; Zordan *et al.* 2007; Lohse *et al.* 2016). Details about this method, the modifications to evaluate white-opaque switching when BSA is the sole nitrogen source, and the modifications to evaluate the effect of BSA on opaque cell stability at 37°C are addressed in the Supplemental Materials and Methods (File S1).

### NanoString Transcriptional Profiling

Transcriptional analysis was conducted using NanoString probes against a set of 59 *C. albicans* genes (one probe per gene). Probes were designed and synthesized by NanoString Technologies. A list of target genes and probe sequences can be found in File S6. Overnight cultures of strains being profiled were grown in SD+AmS+Uri at 25°C, in the morning cells were diluted to a density of 5×10^6^ cells/mL (opaque cells), 2.5×10^6^ cells/mL (white cells), or 1×10^7^ cells/mL (2 hour experiments) in the desired media. Ammonium sulfate samples were harvested eight hours post dilution, other samples were harvested two hours post dilution or when cells were growing logarithmically, as indicated. Cells were pelleted, frozen in liquid nitrogen, and stored at −80°C pending further processing. Total RNA was extracted using a MasterPure™ Yeast RNA Purification Kit (Lucigen MPY03100), the RNA concentration was determined on a Nanodrop 2000C (Thermo Scientific), and RNA quantity was normalized to 400 ng per sample. Samples were processed on a nCounter Sprint Profiler (NanoString Technologies) at the Center for Advanced Technology at the University of California, San Francisco. Read count data was processed using the nSolver analysis software (NanoString Technologies). Two replicates were analyzed for each sample except for wild type opaque cells when BSA was the sole source of nitrogen, in this case four replicates were analyzed.

Raw counts were exported and then analyzed with R (R Core Team 2019) using the tidyverse package (Wickham 2017). All data was normalized based on four housekeeping genes (*PGA59*, MTL*a1, TBP1, HTA1*) whose average expression spans almost three orders of magnitude. The geometric mean of these four housekeeping genes for each sample was calculated, and the difference between the geometric mean of these housekeeping genes across all samples and the geometric mean of these four housekeeping genes in each individual sample was determined. This difference was then subtracted from all reads in that individual sample to normalize the data and these normalized read counts were used for all further comparisons and analyses. Student’s t-tests were performed on log transformed expression data to compare between the ammonium sulfate and BSA conditions for wild type opaque and white cells either logarithmically growing or at the 2hr time point (Figure S3). Significance was evaluated using a 5% false discovery rate (Benjamini-Hochberg procedure). Raw and normalized NanoString transcriptional data can be found in File S6.

### Optical Density Proliferation Assays

The optical density proliferation assays modified our previously reported protocol (Lohse *et al.* 2016) to account for the increased delay before proliferation when BSA was the sole nitrogen source. This method is described in detail in the Supplemental Materials and Methods (File S1).

### Deposited Data and Data Availability

Strains and plasmids are available upon request. All Supplemental Items have been uploaded to the GSA Figshare portal. Supplemental Materials and Methods and Results as well as legends for all supplemental items are included in File S1. A complete list of media, oligonucleotides, plasmids, and strains used in this study can be found in File S2. Additional data for MSP-MS experiments can be found in Figure S1 and File S3. Additional data for proteomics experiments can be found in Table S1 and File S4. Additional data for BSA cleavage assays can be found in Figures S1, S7, and S9. Additional data for cell type switching experiments can be found in Tables S2 and S3. Additional data for flow cytometry based proliferation experiments can be found in Figures S2, S9, S12, S13, S14, and S15 as well as File S5. Additional data for transcriptional profiling experiments can be found in Figures S3, S4, S5, S6, S8, S10, and S11 as well as File S6. Additional data for optical density based proliferation experiments can be found in Figures S7 and S9 as well as File S7.

All raw spectrum (.RAW) files from the MSP-MS and Proteomics experiments reported in this study are available from download from the ProteoSAFe resource (https://proteomics.ucsd.edu/ProteoSAFe/), numbers MSV000085279 (MSP-MS) and MSV000085283 (Proteomics). Reviewer access to these files is available by using login “MSV000085279_reviewer” and password “candidamspms” for the MSP-MS .RAW files and login “MSV000085283_reviewer” and password “candidaprot” for the proteomics files.

## Results

### Opaque cells secrete more proteolytic activity than white cells

To assess secreted proteolytic activity using a global approach, we profiled conditioned media using the Multiplex Substrate Profiling by Mass Spectrometry (MSP-MS) assay (O’Donoghue *et al.* 2012). In brief, white or opaque cell conditioned media were incubated with a library of 228 synthetic 14-mer peptides, and the resulting cleavage products (at 15, 60, and 240 minutes) were identified by liquid chromatography-tandem MS (LC-MS/MS). Opaque cell conditioned media exhibited higher proteolytic activity at each time point examined based on the number of unique cleavage events (377 opaque versus 214 white cleavages, 60 minutes) (Figures 1B and 1C). Other differences in the proteolytic profiles of the two cell types are considered in the Supplemental Results (Figures S1A and S1B, Files S1 and S3). These experiments were performed on conditioned media from cells grown in media containing ammonium sulfate; as such, these results reflect basal, rather than induced, proteolytic activity levels.

We also collected cell-free conditioned media from white and opaque cells, added BSA, and evaluated BSA cleavage over the course of two hours. In agreement with the increased proteolytic activity observed in the MSP-MS assay, opaque cell conditioned media cleaved most of the BSA within 15 minutes and multiple cleavage products were observed by two hours (Figure S1C). White cell conditioned media, on the other hand, cleaved only a small fraction of the BSA at a single position by two hours (Figure S1C).

### Opaque cells secrete different SAPs than white cells

To relate the proteolytic activities of the two cell types to specific proteases, we performed shotgun proteomic analyses on white and opaque cell conditioned media (from cells grown on media containing ammonium sulfate) from three switching-capable strain backgrounds (SC5314, L26, and P37005 (Lockhart *et al.* 2002)) (File S4). Nine unique SAPs were detected in this experiment (Tables 1 and S1). On average, SAPs comprised eight percent of peptides detected in white cell conditioned media and 48 percent of peptides detected in opaque cell conditioned media (Table 1). Consistent with previous transcriptional profiling (Hube *et al.* 1994; White and Agabian 1995), Sap2 was the most abundant SAP secreted by white cells, at four percent of detected peptides (Table 1). For opaque cells, Sap1 was the most abundant SAP, at seventeen percent of detected peptides (Table 1). Sap99 and Sap3 were also abundant in opaque cells, comprising sixteen and eight percent of detected peptides, respectively (Table 1). Also consistent with the previous transcriptional profiling (Hube *et al.* 1994; White and Agabian 1995; Lan *et al.* 2002; Tuch *et al.* 2010; Hernday *et al.* 2013), these three proteases each comprised less than one percent of the peptides detected in white cell conditioned media (Table 1). Likewise, the most abundant white cell SAPs (Sap2 and Sap7) were present at less than 1% of secreted peptides from opaque cells (Table 1). These results show that white and opaque cells secrete two very different spectra of SAPs. Other proteins secreted by each cell type are listed in the Supplemental Results (Files S1 and S4).

**Table 1:** Average total Sap abundance. The total abundance of all Saps detected in white and/or opaque cells averaged across three *C. albicans* strain backgrounds (L26, P37005, SC5314). Sap4, Sap5, and Sap6, which are normally associated with biofilms, were not detected in any sample of either cell type.

| Protein | Average Percent of Total Counts, All White Strains | St. Dev. Percent of Total Counts, All White Strains | Average Percent of Total Counts, All Opaque Strains | St. Dev. Percent of Total Counts, All Opaque Strains |
| --- | --- | --- | --- | --- |
| Sap1 | 0.42 | 0.03 | 17.36 | 2.21 |
| Sap2 | 3.61 | 2.07 | 0.49 | 0.23 |
| Sap3 | 0.51 | 0.23 | 8.37 | 0.64 |
| Sap7 | 1.85 | 0.55 | 0.00 | 0.00 |
| Sap8 | 0.14 | 0.10 | 0.82 | 0.59 |
| Sap9 | 0.50 | 0.06 | 0.04 | 0.05 |
| Sap10 | 0.85 | 0.27 | 0.26 | 0.03 |
| Sap98 | 0.00 | 0.00 | 4.21 | 2.57 |
| Sap99 | 0.57 | 0.18 | 16.09 | 6.33 |
| All Saps | 8.44 | 2.58 | 47.64 | 9.90 |
| The total abundance of all Saps detected in white and/or opaque cells averaged across three *C. albicans* strain backgrounds (L26, P37005, SC5314). Sap4, Sap5, and Sap6, which are normally associated with biofilms, were not detected in any sample of either cell type. |  |  |  |  |

### Opaque cells are more efficient at using proteins as a sole nitrogen source

Soll and colleagues previously reported that opaque cells from the WO-1 strain proliferate sooner (that is, there is a shorter delay after inoculation before they begin logarithmic proliferation) than white cells when BSA is the sole nitrogen source (Srikantha *et al.* 1995; Kvaal *et al.* 1999). Consistent with this result, we found that opaque cells from the three strain backgrounds used in this study began logarithmic growth sooner than white cells when BSA was the only source of nitrogen (roughly 18-24 versus 72-120 hours, depending on starting cell density and nutrient carry over from dilutions) (Figures 2A, 6B, and S9B). We also observed that opaque cells reach a slightly higher maximum proliferation rate, doubling in slightly less than four hours as opposed to every five hours for white cells (File S5). The presence of BSA does not appear to increase white-opaque switching in either direction and does not block temperature (37°C) induced *en masse* opaque-to-white switching (SI Results, Figures S2A and S2B, Table S2).

To determine whether similar proliferation differences were observed with proteins other than BSA, we measured proliferation on human serum albumin (HSA), hemoglobin, and myoglobin. Opaque cells began proliferating at 30-40 hours when HSA was the nitrogen source compared with no growth after 90 hours for white cells (Figure 2B). Opaque cells also began proliferating sooner and slightly faster than white cells when hemoglobin was the nitrogen source, although the difference between the two cell types was less pronounced (proliferation after 8-16 versus 20-40 hours, doubling roughly every two as opposed to three hours) (Figure 2B, File S5). In contrast to the other protein sources, myoglobin seemed equally well utilized by white and opaque cells as both cell types rapidly began to proliferate, although opaque cells still proliferated slightly faster (doubling every 3.5 versus five hours) (Figure 2B, File S5). These results suggest that opaque cells are generally more efficient than white cells at utilizing proteins as a nitrogen source, but that protein-specific characteristics (e.g. primary sequence and/or folding) affect the difference with which the two cell types can utilize specific proteins.

*C. albicans* is unusual among fungi in that it can utilize free amino acids as a carbon source in addition to their more common utilization as a nitrogen source (Vylkova *et al.* 2011; Priest and Lorenz 2015; Ene *et al.* 2016). To determine whether proteins could be used as a sole carbon source, we examined the proliferation of both cell types on media containing BSA but lacking glucose (both with and without ammonium sulfate). Neither white nor opaque cells reached their typical final cell density (roughly 10^8^ cell / mL) in these conditions (Figure S2C), indicating that *C. albicans*, although it can utilize proteins as a nitrogen source, cannot efficiently use proteins, at least across the scope of this experiment, as their sole carbon source.

### White and Opaque cells express different *SAP*s in response to nitrogen source

To understand the response of white and opaque cells to different nitrogen sources, we used NanoString probes to measure expression of the *SAP* family genes and 13 transporters (8 *OPT*s, 2 *PTR*s, 2 ammonium permeases, and 1 amino acid permease) on ammonium sulfate, BSA, myoglobin, and hemoglobin, as sole nitrogen sources, as well as nitrogen-depleted medium (Figure 3A). To allow for an equivalent comparison between cell types on BSA, we profiled both cell types on BSA (1) after two hours in the new medium and (2) when they began logarithmic proliferation (∼24 hours for opaque cells, ∼96 hours for white cells).

We first consider proliferation on ammonium sulfate, which we consider to be the “basal” (i.e. nitrogen-replete) expression program. Consistent with previous reports (Hube *et al.* 1994; White and Agabian 1995; Lan *et al.* 2002; Tuch *et al.* 2010; Hernday *et al.* 2013), white cells (relative to opaque cells on the same medium) express higher levels of the oligopeptide transporters *OPT1* (8-fold), *OPT3* (40-fold), *OPT7* (14-fold), and *PTR22* (6-fold) (Figure 3B, File S6). Although Sap2 was detected in white cell conditioned media in the proteomics experiments discussed above, *SAP2* transcript levels were similarly low in both white and opaque cells growing on ammonium sulfate. In agreement with the proteomics experiments, opaque cells, on the other hand, express higher levels of *SAP1* (160-fold), *SAP3* (5.9-fold), *SAP99* (15-fold), and the oligopeptide transporter *OPT4* (65-fold) relative to white cells (Figure 3B, File S6).

In contrast to the “basal” expression profiles observed in ammonium sulfate, the induced response is seen when BSA is provided as the sole nitrogen source. On such media, we observe (1) groups of genes that are upregulated in both cell types (although these are often up-regulated to a greater extent in opaque cells), including *SAP2* (1450-fold in white cells, 420-fold in opaque cells), *SAP3* (18-fold in white cells, 630-fold in opaque cells), *OPT2* (70-fold in white cells, 210-fold in opaque cells), *OPT5* (170-fold in white cells, 230-fold in opaque cells), and *PTR2* (120-fold in white cells, 580-fold in opaque cells), (2) genes that are only up-regulated in white cells such as *OPT1* (15-fold), and (3) genes that are only up-regulated in opaque cells such as *SAP8* (900-fold), *OPT7* (19-fold), and *SAP1* (5-fold) (Figures 3B, 3C, S3, File S6). Both cell types’ responses to BSA as the sole nitrogen source are similar to those when either hemoglobin or myoglobin are the sole nitrogen source (Figure S4, File S6).

We also observed the transcriptional response of white and opaque cells before they had a chance to adapt to proteins as their sole nitrogen source and proliferate (2 hours post-inoculation) and to simple nitrogen starvation (SI Results, Figures S5 and S6, File S6). These results suggest that both cell types initially respond to nitrogen starvation (in much the same way) but eventually this response dissipates, and the SAP-transporter program, which differs markedly between white and opaque cells, takes over (Figure 3D).

Several general points emerge from this transcriptional analysis. First, white and opaque cells differ in their basal expression of the *SAP*s and *OPT*s, that is, their expression of these genes in nitrogen replete medium (Figure 3D). Superimposed on these distinct basal expression patterns are the induction of different sets of *SAP*s and *OPT*s in response to the presence of protein as the sole nitrogen source. In opaque cells, some of the basal opaque-specific genes are further-up-regulated and a series of new genes are transcribed, giving an induced expression pattern that is markedly different from that of white cells (Figure 3D).

### Contributions of individual SAPs

The aspartyl protease inhibitor Pepstatin A blocked both basal opaque BSA cleavage (Figure S7A) and opaque cell proliferation on BSA as the sole nitrogen source at concentrations that did not affect proliferation on ammonium sulfate (Figure S7B). These results indicate that both phenotypes are dependent on *SAP* protease activity. To identify the specific *SAP*(s) responsible, we examined basal opaque BSA cleavage and opaque cell proliferation on BSA in a set of strains where combinations of either four or five of the opaque-expressed *SAP*s (*SAP1, SAP2, SAP3, SAP8*, and *SAP99*) were deleted. We found that basal opaque BSA cleavage was abolished when all five of these *SAP*s were deleted (Figure S7C). When four of the five SAPs were deleted, we found that the presence of either *SAP1* or *SAP3* could maintain BSA cleavage at near wild type levels but that *SAP2, SAP8*, or *SAP99* alone did not support basal opaque BSA cleavage (Figure S7C). Efficient opaque cell proliferation on BSA as a sole nitrogen source was abolished (for at least 7 days, the extent of the experiment) when all five *SAP*s were deleted (Figure 4). The presence of either *SAP1, SAP2, SAP3*, or *SAP8*, but not *SAP99*, was sufficient to allow opaque cell proliferation on BSA, although this proliferation was delayed relative to the wild type strain (Figure 4, File S5). Thus, “basal” cleavage by opaque cells depends on *SAP1* and *SAP3*, while efficient proliferation on BSA as a sole nitrogen source can be sustained by any of the four highly induced *SAP*s (*SAP1, SAP2, SAP3*, and *SAP8*). We did not observe significant changes in the basal expression of *SAP4*-*SAP7, SAP9, SAP10*, or *SAP30* when *SAP1, SAP2, SAP3, SAP8*, and *SAP99* were deleted (Figure S8, File S6), suggesting that there is no compensatory up-regulation in response to the loss of basal opaque cell *SAP* expression. The only exception was an increase in *SAP98* expression when *SAP99* was deleted, but these two genes are adjacent and we believe that this upregulation is likely the result of the nearby genetic manipulation. We also note that deletions of either four or five of the opaque-expressed *SAP*s (*SAP1, SAP2, SAP3, SAP8*, and *SAP99*) do not appear to block or increase white-opaque switching in either direction (Table S3, SI Results, File S1).

### Identification of transcriptional regulators needed for the response of opaque cells to protein as the sole nitrogen source

To identify regulators controlling opaque cells’ basal proteolytic activity and use of proteins as a nitrogen source (which requires the induced response), we screened a library of 188 opaque transcription regulator deletion strains (Lohse *et al.* 2016) for (1) loss of BSA cleavage in conditioned media and (2) limited proliferation when BSA was the sole nitrogen source. Only two of the 188 opaque strains tested, the *efg1* and *wor3* deletions, lacked basal opaque cell proteolytic activity (Figure S9A). Both Efg1 and Wor3 are known regulators of white-opaque switching and both are required for full expression of the opaque cell transcriptional program (Sonneborn *et al.* 1999; Zordan *et al.* 2007; Hernday *et al.* 2013; Lohse *et al.* 2013). Efg1 has previously been linked to *SAP* expression in the oral epithelial cell and parenchymal organ invasion models, although these experiments were conducted with white cells (Felk *et al.* 2002; Korting *et al.* 2003). Because *WOR3* expression in opaque cells is dependent on the presence of Efg1 (Hernday *et al.* 2013), the *efg1* phenotype probably reflects a loss of *WOR3* expression. Consistent with this idea, expression of *SAP1, SAP3, SAP99*, and *OPT4* was lost in both opaque *wor3* and *efg1* deletion strains grown on ammonium sulfate (Figure 5).

In the second screen (failure of opaque cells to proliferate on BSA as the sole nitrogen source), the *stp1* deletion strain showed a pronounced proliferation defect, and the *csr1* deletion strain exhibited a milder defect (Figure S9B, File S1, File S7). Stp1 was previously shown to be necessary for white cell proliferation on BSA (Martínez and Ljungdahl 2005). Neither Stp1 nor Csr1 were needed for opaque cell basal opaque proteolytic activity (SI Results, Figures 5 and S9A, File S1).

To understand the relationship between Stp1, Csr1, Efg1, and Wor3 in the induced opaque-cell response, we took advantage of the constitutively active allele of Stp1, Stp1 Δ2-61 (Martínez and Ljungdahl 2005). When Stp1 is constitutively activated in this way, both white and opaque cells exhibit gene expression in ammonium sulfate patterns similar to those seen in wild type cells proliferating on proteins as a sole nitrogen source (Figures 5 and S10, File S1). Conversely, neither the SAPs nor the transporters are up-regulated in opaque cells by proteins as a sole nitrogen source when *STP1* is deleted (Figure S11A, Files S1 and S6). Additional experiments showed that Wor3 and Efg1 are needed for both the basal and induced responses in opaque cells while Csr1 is needed only for the induced response (Figures 5 and S10, Files S1 and S6). We conclude from these experiments that activation of Stp1 is the key initiating event in both the white and opaque cell responses to protein as the sole nitrogen source, but that Wor3, Efg1, and Csr1 are needed for the full elaboration of the response in opaque cells (Figure 5B).

Stp1 activation in white cells is dependent on the membrane bound sensor Ssy1 (Csy1); deletion of *SSY1* in white cells blocks Stp1 truncation, and thus activation, in response to extracellular amino acids (Martínez and Ljungdahl 2005). We deleted *SSY1* to determine if it was also necessary for the opaque cell induced response. Although proliferation of an opaque *ssy1* deletion strain on BSA as the sole nitrogen source was slightly delayed (roughly 24-36 hours versus 18-24 hours) and slower (doubling every 4.5 versus four hours) than wild type (Figure S9B, File S5), the transcriptional profile of opaque *ssy1* deletion cells growing logarithmically on BSA was similar to that of wild type opaque cells (Figure S11, File S6). Thus, we conclude that the protein-induced program in opaque cells requires Stp1, Wor3, Efg1, and Csr1, while the protein-induced program in white cells requires Stp1 and Ssy1 only. These results are summarized in Figure 5B.

### A minority opaque cell population supports white cell proliferation on BSA as a sole nitrogen source

To test whether the induction of the *SAP*s in opaque cells could help white cells utilize BSA as a sole nitrogen source, we co-incubated white and opaque cells tagged with different fluorescent markers and tracked the proliferation of each cell type over several days using flow cytometry (Figure 6A). In the presence of BSA as the sole nitrogen source, pure opaque cell and white cell populations began logarithmic proliferation after approximately 24 and 60-72 hours, respectively, with the proliferation rate, once the cells begin to divide, being faster in opaque cells (doubling every 3.5-4.5 hours versus every 6-9 hours) (Figure 6B, File S5). These results are consistent with those described earlier (Fig. 2A), indicating that the fluorescent markers did not significantly affect the relative growth rates. In populations that contained 20% opaque cells and 80% white cells, both cell types began proliferation after 24-36 hours and both proliferated at more similar rates (Figure 6B, File S5). Even a small fraction of opaque cells (<5%) had a significant effect on white cell proliferation in the presence of BSA as the sole nitrogen source, and we found that the maximum proliferation rate of white cells increased with the fraction of opaque cells (Figure 6B, File S5). To further explore this “helping” effect, we co-cultured wild type opaque cells with the opaque quintuple *SAP* deletion strain and with the opaque *STP1* deletion strain (neither of which can efficiently utilize BSA as the sole nitrogen source) and found that wild type opaque cells could completely rescue the quintuple *SAP* deletion strain (which actually proliferated slightly faster than wild type, doubling every 2.9 versus 3.2 hours) and could partially rescue the *stp1* deletion strain (which doubled every 5.8 hours) (Figure S12, File S5). These observations suggest that the growth defect of the quintuple *SAP* deletion stems solely from a lack of SAP-cleaved peptides, but the growth defect of the *stp1* deletion strain also includes a failure to take up cleavage products due to reduced expression of oligopeptide transporters.

Given that wild type white cells responded better than the opaque *STP1* deletion strain did in these co-culture experiments, we hypothesized that white cells mounted a specific response to BSA as a sole nitrogen source but only in the presence of opaque cells. To test this idea, we performed co-culture experiments with a GFP tagged opaque strain and white strains with mCherry reporters driven by the *OPT1, OPT2, SAP2*, or *UGA4* promoters. In pure populations of white cells exposed to BSA as the sole nitrogen source, neither *SAP2* nor *OPT1* expression is fully up-regulated within 60 hours and, conversely, *OPT2* and *UGA4* (which are induced simply by nitrogen starvation) are up-regulated within 18 hours and remain fully induced after 60 hours (Figures 6C and S13). In the presence of opaque cells, however, there is a noticeable increase in *SAP2* expression after 19 hours which continues to increase until it is fully expressed in all white cells by 42 hours (Figure 6C). Likewise, *OPT1* levels begin to increase (relative to a pure white cell population) within 18 hours and appear fully induced within 24 hours (Figure S13). Expression of *OPT2* and *UGA4*, on the other hand, noticeably begins to decrease (relative to a pure white cell population) after 18-24 hours and reached their minimum within 42 hours in the presence of opaque cells (Figure S13). These results indicate that not only does the presence of opaque cells allow white cells to proliferate in the presence of BSA as the sole nitrogen source, but that the white cells respond differently (sooner) to BSA as the sole nitrogen source in the presence of opaque cells. The white cell response appears to be due to the opaque cells’ ability to cleave BSA, rather than a specific signal from them: we observe a similar response when white cells were provided with BSA that had been pretreated with Proteinase K (Figure S14).

To test whether the helping effect of opaque cells for white cells extended across species, we performed similar co-culture experiments with *C. dubliniensis, C. tropicalis*, and *C. parapsilosis*, none of which can efficiently utilize BSA as the sole nitrogen source (Figure S15). All three of these species showed significantly improved proliferation when co-cultured with wild type opaque *C. albicans* (Figure S15), but not with wild type white *C. albicans* cells (Figure S15).

## Discussion

*Candida albicans*, a fungal component of the human microbiome and an opportunistic pathogen of humans, can switch between two heritable states, called white and opaque. In this paper, we identify a highly inducible gene expression program that is unique to opaque cells, enabling them to efficiently utilize proteins as a sole source of nitrogen. We demonstrate, using proteomics, multiplex protease substrate profiling, and transcriptional analysis, that a series of secreted aspartyl proteases (SAPs) are constitutively secreted by opaque cells, comprising almost half of the secreted proteins. When protein is present as the sole nitrogen source, the *SAP*s (already expressed at high levels) are up-regulated even further, along with peptide transporters. We show that the protein-induced response is not simply an up-regulation of the basal pattern: different *SAP*s and transporters are up-regulated to produce an entirely new pattern of gene expression. We show that these secreted proteases degrade generic proteins, whose peptides can then be used as a nitrogen source. As a result of this opaque-cell specific response, opaque cells begin proliferating much sooner than white cells in media where protein is the sole nitrogen source; in nearly all other types of media, white cells outcompete opaque cells. Using deletion mutants, we identified the individual *SAP*s required for opaque cells to efficiently utilize protein as a nitrogen source. In contrast to white cells, where a single *SAP* (*SAP2*) allows limited growth under these conditions (Hube *et al.* 1997), up-regulation of any one of four *SAP*s can suffice for efficient proliferation of opaque cells. Activation of Stp1, a previously identified regulator of *SAP* genes in white cells (Martínez and Ljungdahl 2005), is also key for the opaque-specific up-regulation, although the response differs greatly between white and opaque cells. Compared to white cells, opaque cells appear to activate Stp1 in a different way and have incorporated at least three additional regulators (Wor3, Efg1, and Csr1) that act in concert with Stp1. It seems likely that these differences contribute to both the timing of the response in opaque cells as well as its magnitude.

White and opaque cells differ in many properties and are often thought of as being specialized for different niches in their mammalian hosts. In this paper, we show that, in mixed cultures of white and opaque cells, a minority of opaque cells, by providing massive amounts of secreted proteases, can benefit the entire culture. This result suggests that, rather than occupying separate niches in the host, the two types of cells can work together to produce a population that is robust to changing environments. White and opaque cells carry identical genomes and it is an intriguing idea that *C. albicans’* ability to adapt to a wide range of environments might be achieved, at least in part, by partitioning expression of the genome between two different types of cooperating cells, where each cell type has specific, but complementary, metabolic characteristics. Such a partnership need not be limited to nitrogen acquisition, it could extend to a wide-range of other *SAP*-mediated host interactions (e.g. interactions with bacteria (Dutton *et al.* 2016), adhesion to host cells (Watts *et al.* 1998; Bektic *et al.* 2001; Albrecht *et al.* 2006), protection from host defense proteins (Borg-von Zepelin *et al.* 1998; Gropp *et al.* 2009; Meiller *et al.* 2009; Rapala-Kozik *et al.* 2010, 2015; Bochenska *et al.* 2015; Kozik *et al.* 2015), activation of immune responses (Schaller *et al.* 2005; Hornbach *et al.* 2009; Pietrella *et al.* 2010; Pericolini *et al.* 2015; Gabrielli *et al.* 2016)) as well as additional differences between the two cell types that extend beyond the secreted aspartyl proteases.

Finally, we note that the benefits of *SAP*-mediated nitrogen acquisition by opaque cells are not limited to *C. albicans*. We showed that three other species (*C. dubliniensis, C. tropicalis*, and *C. parapsilosis*), each of which, on its own, cannot efficiently proliferate on BSA as a sole nitrogen source, can do so in the presence of *C. albicans* opaque cells. There is no *a priori* reason that more distantly related fungi, or even bacteria, with the capacity to uptake and process short peptides could not also benefit from the presence of *C. albicans* opaque cells.

## Acknowledgements

We thank Haley Gause, Carrie Graham, and Niyati Rodricks for advice. We thank Ananda Mendoza for technical support. Support was provided by the UCSF Mass Spectrometry Facility (A.L. Burlingame, Director) supported by the Dr. Miriam And Sheldon G. Adelson Medical Research Foundation (AMRF) and the National Institutes of Health (NIH) – NIGMS NIH P41GM103481. This work was supported by NIH grants F32CA168150 (to M.B.W), P50AI150476 (to C.S.C) and R01AI049187 (to A.D.J.). This investigation has also been aided by a grant from The Jane Coffin Childs Memorial Fund for Medical Research (to N.Z.). The content is the sole responsibility of the authors and does not represent the views of the NIH or other funding agencies. Neither the NIH nor other funding agencies had any role in the design of the study, in the collection, analyses, or interpretation of data, in the writing of the manuscript, or in the decision to publish the results.

## Conflict of Interest Statement

Alexander D. Johnson is a cofounder of BioSynesis, Inc., a company developing inhibitors and diagnostics of *C. albicans* biofilms. Matthew Lohse was formerly an employee and currently is a consultant for BioSynesis, Inc.

